# Development of an insect sample fractionizer for biodiversity research

**DOI:** 10.1101/2022.11.04.515206

**Authors:** Thomas Hörren, Martin Sorg, Caspar A. Hallmann, Werner Stenmans, Axel Ssymank, Hannes Theumert, Jan Jürgens, Bernhard Fleischer, Heinz Schwan

## Abstract

We describe a new mechanical tool for dividing mixed insects and other invertebrate samples into subsamples. The device enables the division to equal parts by means of a movable hemispherical bowl and a separating disc. Due to the complete stainless steel manufacturing, the sample divider is sterilizable by using chemicals or heating and thus suitable for DNA-based methods. The production of equally sized subsamples is of particular importance for biodiversity studies today, especially when using metabarcoding combined with insect homogenisation for species determination of mixed insect samples. The device allows sub-samples to be analyzed separately using the same or different methods, or getting archived for museal preservation and future research.

## Introduction

Biodiversity research in terrestrial ecosystems needs a holistic overview of its components that characterizes and interacts with a specific habitat. Insects in particular represent the vast majority of the diversity of any terrestrial biotope in most climate zones. These biotopes would simply not be imaginable without their diverse, regulating functions and influences. Because of this high diversity of often several thousand species at a given point of investigation (1), their total species and abundance composition also contains by far the most precise source of information about the local state of nature and the local processes of biodiversity change. Efficient insect trapping techniques such as malaise traps (2) or gressitt traps (3) and similar methods are able to capture large parts of this insect composition acting at a survey point in their samples. Modern species identification methods such as metabarcoding are in principle capable of analyzing the species diversity of these mixed insect samples. However, neither is the entire insect diversity of certain countries taxonomically described, nor are the barcodes of all species comprehensively known. When using destructive laboratory methods for the purpose of metabarcoding via homogenization of mixed species samples, this means the loss of processing and preservation potential for a wide variety of purposes. It is therefore advisable to produce subsamples for a wide variety of purposes and not always to use complete samples for certain laboratory processes to clarify specific questions.

## Development and components

The insect sample fractionizer was developed by the Entomological Society Krefeld in cooperation with our coauthors from a technical school for mechanical engineering and a specialist steel construction company over a period of more than one year. After first successful experiments in 2018 with plastic parts, the development of two previous stainless steel prototypes during 2019 was necessary to achieve the final model described here.

### Characteristics, material and additional equipment

Base plate with welded supports that carry a semi-hollow divider hemisphere. Four threaded screws (Imbus) for leveling the base plate. Semi-hollow sphere, D = 300mm, custom-made, polished inside. Axle, welded, carries the semi-hollow sphere. Locking pin, screwed on. Locking disc, welded. Two outlet funnels and divider plate, welded on. Whole construction made of stainless steel type 1.4301. The total weight measures 13.5kg. The upper part (semi-hollow sphere) measures 5.8 kg. Detailed descriptions and measurements see supplement 1-2.

Equipment required in addition: Spirit level. Precision scale with a range of 0-500g, accuracy of 0.1g. Allen key, size 2.5 and 6. Squirt bottle suitable for ethyl alcohol/ethanol/EtOH. Collecting vessels (e.g. beakers, heat-resistant laboratory glass, 1000ml - high form). Rounded, heat-resistant metal or glass rods. Sieves made of stainless steel for the determination of the ethanol-moist biomass of the mixed insect samples. Indexing bolts, also known as locking bolts, are used in connection with e.g. welded-on indexing discs. These are structural components that allow moving components to be adjusted quickly and in a defined manner. By using the pull button or the pull ring, a pin is moved out of the counterpart in order to bring components into the appropriate position and fix them.

**Fig. 1.**
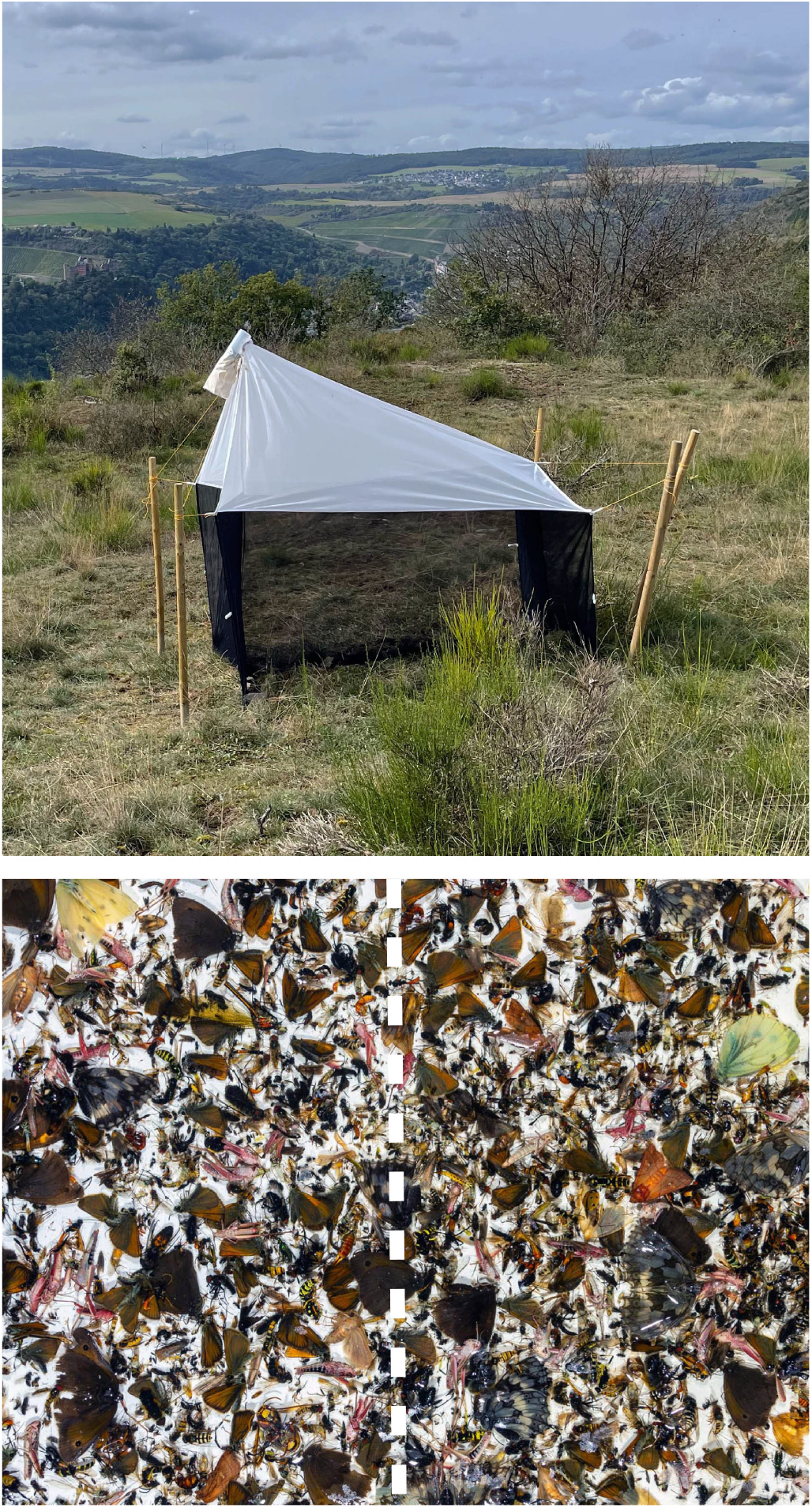
A malaise trap of the Entomological Society Krefeld (4)(5) (top) and a typical weekly early summer catch result with a drawn line to symbolize the separation of these diverse sample types into two subsamples (below).

**Fig. 2.**
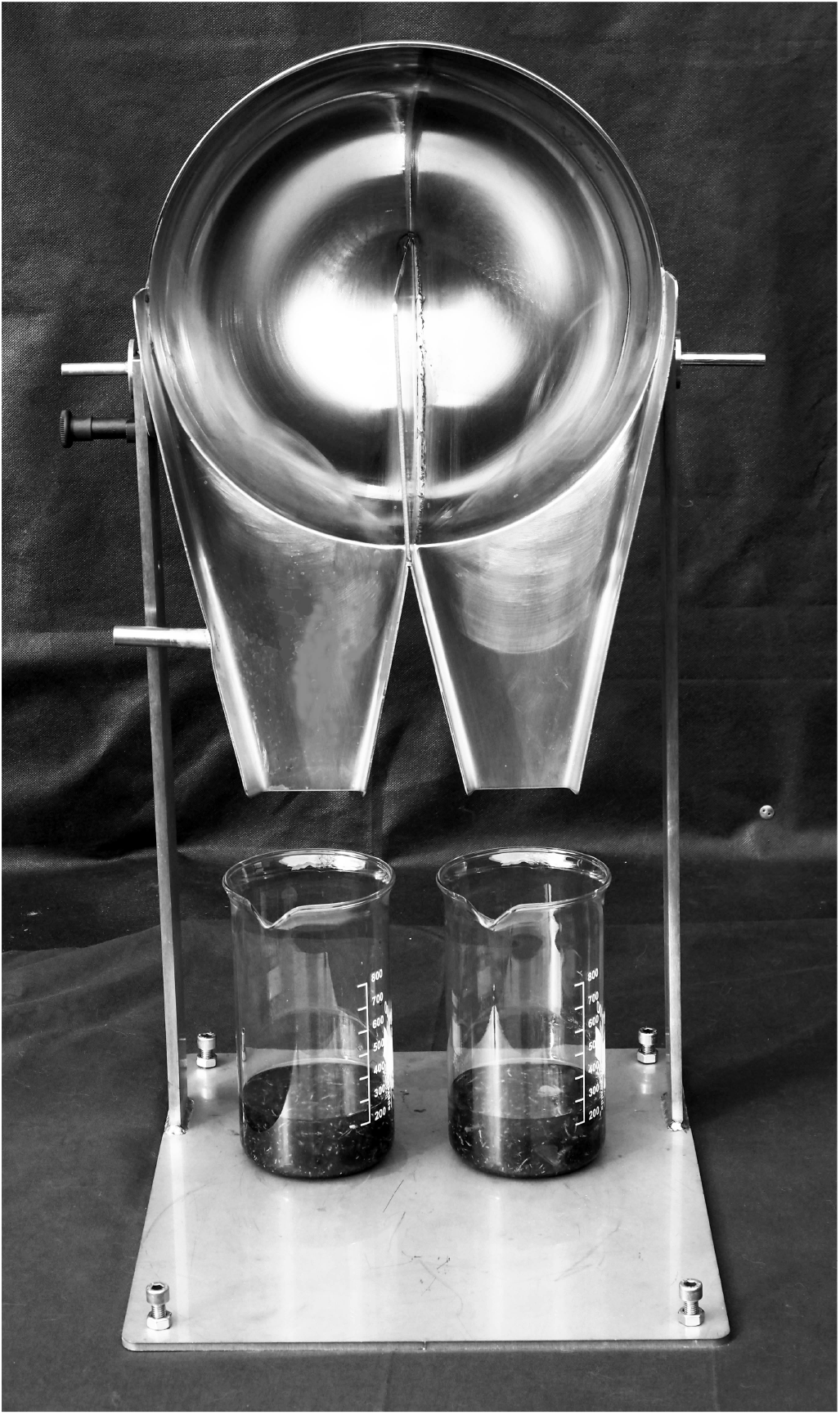
The sample fractionizer with the locking bolt for fixing the position of the hemispherical dish (to the left of the hemispherical dish) and its 4 height-adjustable screws in the corners of the base plate for leveling.

**Fig. 3.**
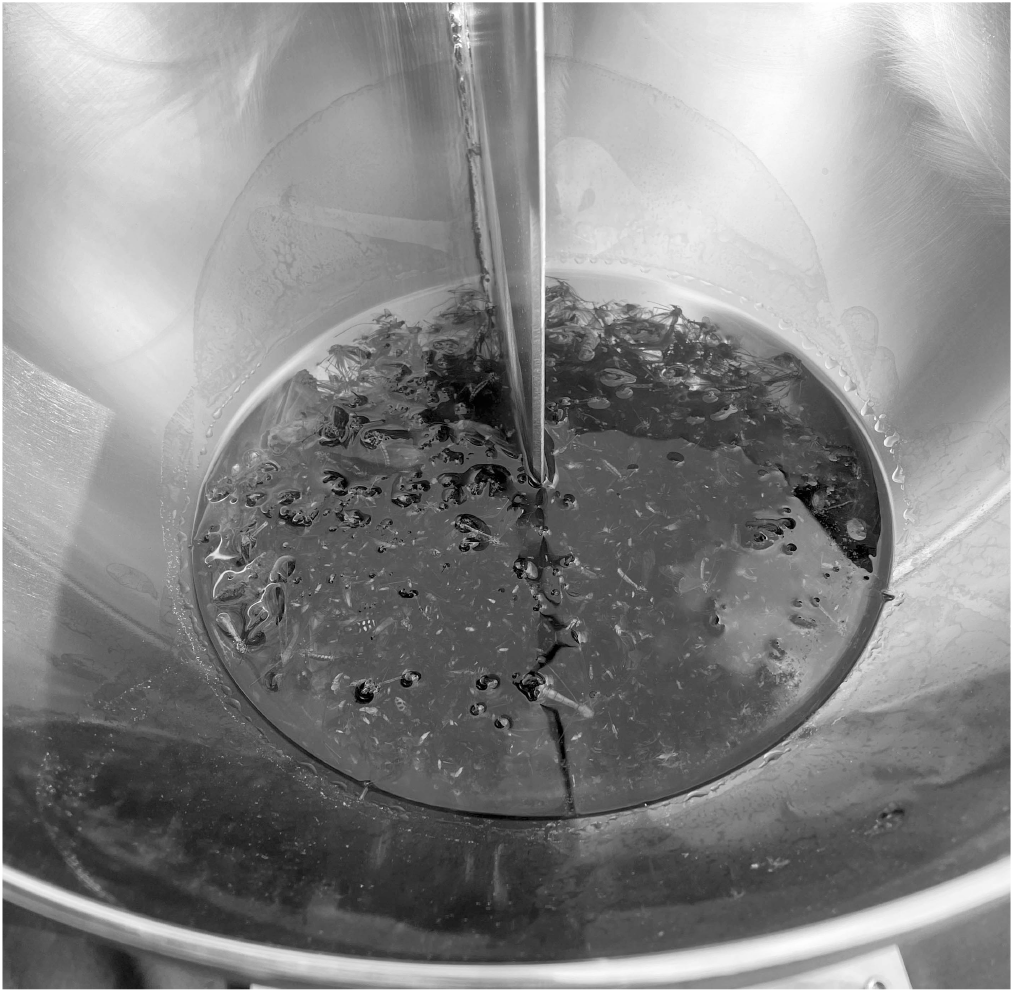
The insect sample fractionizer in action. Axial rotation distributes the invertebrates in the total volume into equally sized parts.

## Recommendations for use

The sample fractionizer was developed especially for biomass-rich (5), high diversity malaise trap samples that needed to be processed DNA-sterile. We recommend to use the sample fractionizer for invertebrate samples preserved in ethanol of high concentration.

### Application description

Before the start of each new workflow, the sample fractionizer should be precisely adjusted on the work surface.

For calibration, about 500g of water is weighed out and placed in the rear part of the half-hollow sphere. Two beakers, each with a capacity of 1000ml and a known tare weight, are passed over the two spouts that are to hold the divided sample quantity. The sample fractionizer is now moved forward and downward with a consistentmovement until the drainages are in a vertical position. This position is locked so that the fractionizes water sample can drip completely into the beakers. When the dripping process stops, the weight of both beakers is determined. Subtracting the tare, the divided amount of water shows the precision of the fractionation process for a liquid. A positive result exists if the difference between the two partial quantities is max. 1 gram. If the difference is greater, the set-up should be readjusted.

The amount of insects filled in should be covered by alcohol in a way that the insects easy move around. For malaise trap samples a maximum insect quantity of up to approx. 50g can be used for one dividing process. The capacity maximum amount of insects and alcohol using this device is close to 500ml.

More hints for the separation process:

No contact with the built-in separating plate or e.g. the filled quantity should not touch the separating plate at the beginning of the separating process. The separating plate protrudes 10mm beyond the center of the half hollow sphere, which ensures that the 1:1 separated sample is not mixed again in the rear edge area. The sample divider is free to move forwards and backwards around the axis. In comparison to alcohol, insects behave like an inert mass during the dividing process. By slowly increasing back and forth movement, the alcohol including the insects becomes freely movable. Once such a state is reached, the separation process is carried out by moving the dividing trays downward and forward evenly but decisively. The dividing chutes are locked in a vertical position. The separation result can be as follows:

a. Ideally, the mixture of insects and alcohol is completely divided 1:1 and is available in 2 beakers of 1000ml each.
b. Insects have partly remained in the edge area of the divider and are rinsed into the beakers with alcohol from the spray bottle.
c. Moving too quickly can cause individual insects to get caught on the metal edges in the outer areas and not be split properly..
d. Moving too slowly, leaves possibly many insects in the sample fractionizer. The alcohol consumption is considerably increased by necessary extensive rinsing.

With vertically downward fixed outlets, the sample divider must be rinsed well with help of ethanol in the spray bottle.

### Sterilizing of the insect sample fractionizer

After completion of each working procedure, the sample fractionizer is cleaned out with a damp cloth to remove residues such as butterfly scales, these would otherwise burn in as residues if heating is used for the sterilization process. The insect sample fractionizer can be cleaned with sterilizing chemicals. Since the device is made of stainless steel, e. g. a sterilization process after cleaning with ethanol can also be carried out using high temperatures, which prevents the unwanted transfer of DNA traces between samples. This can be done using a gas burner or by igniting a liquid film of ethanol that has been sprayed on. The interior of the divider is for this porpose sprayed with a few strokes of a spray bottle containing high concentration ethanol and the alcohol is lit with a long-handled lighter. The same methodology can be used for the accompanying tools, which are also made of stainless steel or heat-resistant laboratory glass. In order to clearly illustrate the process, short instructional films were produced (see supplement 3).

### Safety instructions for sterilization of working equipment by burning ethanol

Do not use in closed rooms and outside appropriately equipped fume hoods. When used outdoors, no flammable material must be accessible in the vicinity. Depending on the amount of ethanol, burning can least for up to several minutes with a flame, e.g. between 30 and 50cm high, which can easily ignite combustible materials in the surrounding area due to crosswinds. The process must be supervised until all flames are completely extinguished. A fire extinguisher should be readily available within reach, and more extensive fire protection measures should be taken. Wear protective goggles when working with ethanol and notice safety data sheets. Never add ethanol to the burning fire or the still heated working tools, this could lead to dangerous deflagrations. At high air temperatures of e.g. over 30 degrees Celsius, slight deflagration will also occur during ignition. We recommend that the executing person should have training and experience in proper laboratory practice for the above-mentioned processes (e.g. chemical laboratory technician) and of course should be familiar with the associated fire protection measures, accident prevention regulations and safety data sheets. In principle, the institutional requirements at the point of execution must be observed. For the sterilization process of the equipment see video-links under supplement 3

## Conclusions

The insect sample fractionizer is the first tool that enables for a standardized sample fraction of mixed insect samples, e.g. from the results of biodiversity studies using efficient trap technology. Due to the complete stainless steel manufacturing, the sample fractionizer is sterilizable by chemicals or heating and thus suitable for DNA-based methods for species or operational taxa analyses. The invertebrate sample fractionizer can be used for a variety of research issues in biodiversity research. The design features allow for a splitting of an insect composite sample whose closeness to the equal splitting for the insect biomass can be checked by determining the wet biomass of the subsamples. Of course, species that are randomly distributed in only one or very few individuals before the fractionizing process (so called singletons) can only be present in one of the subsamples. These patterns correspond to the natural distribution pattern within ecological societies of mixed samples as well as the basic ecological features of population ecology. In the case of e.g. malaise traps, however, there are higher numbers of individuals for species that are represented in a habitat with a corresponding activity abundance. If questions of a more complete recording of species with low abundances are in the foreground of an investigation, then of course a decision should be made which sample sizes are required. In these cases, of course, the question arises as to how many trap units per area size are sufficient or more efficient trap techniques are required.

## ACKNOWLEDGEMENTS

Conceptual framework and development of methodologies connected with malaise trap samples of the EVK (TH, MS, HS, WS) was funded by the German Federal Ministry for the Environment, Nature Conservation and Nuclear Safety (BMU), handled by the The German Federal Agency for Nature Conservation (BfN), grant number FKZ 3516850400. The manufaction of prototypes of the insect sample fractionizer was funded by the Ministry for Environment, Agriculture, Conservation and Consumer Protection of the German State of North Rhine-Westphalia (MULNV) (No. III-1-620.08). We would also like to thank many experts from the Entomological Society Krefeld for advice and assistance during the development of the new tool.

## Author contributions

TH, MS, HS and WS developed the conceptual framework. HS and MS realized initial concepts in models. HS, JJ, BF and HT designed and produced the prototypes. TH and MS drafted the first version of the manuscript. All authors contributed to the article and approved the version to be published.

**Supplement 1**

Technical drawings of the sample fractionizer, .pdf file: http://entomologica.org/tools/ISF01.pdf

**Supplement 2**

Full resolution 3D-CAD-model of the sample fractionizer, .pdf file: http://entomologica.org/tools/ISF02.pdf

**Supplement 3**

Video documentation of the sterilization process of the equipment:

https://youtu.be/bSo6el6yY-8

https://youtu.be/RVdPEBPi0y8

https://youtu.be/kA09Mhm2cmw

## Notes

https://youtu.be/RVdPEBPi0y8

https://youtu.be/kA09Mhm2cmw

### Competing Interest Statement

The authors have declared no competing interest.

